# Estimating sampling completeness of interactions in quantitative bipartite ecological networks: incorporating variation in species’ specialisation

**DOI:** 10.1101/195917

**Authors:** Callum J. Macgregor, Darren M. Evans, Michael J.O. Pocock

## Abstract

**Background:** The analysis of ecological networks can be affected by sampling effort, potentially leading to bias. Ecological network structure is often summarised by descriptive metrics but these metrics can vary according to the proportion of the total interactions that have been observed. Therefore, to know the likely degree of bias, it is valuable to estimate the total number of interactions in a network, and so calculate the proportion of interactions that have been observed (sampling completeness of interactions). Existing approaches to estimate sampling completeness of interactions use the Chao family of asymptotic species richness estimators to predict the total number of interactions, but do not fully utilise information about the relative specialisation of species within the network.

**Results:** Here, we propose a modification of previously-used methods, that places equal weight on each interaction (whether or not it has been observed), rather than on each species. Our approach is therefore equivalent to weighting the interaction sampling completeness of each species in the network according to its relative specialisation. We demonstrate that, for the subset of species that are observed and when assuming that species richness estimators accurately project the number of unobserved interactions per observed species, our approach is mathematically more accurate. Our approach can be universally applied to any quantitative, bipartite network.

We propose two methods to estimation using our approach, using abundance-based and incidence-based species richness estimators respectively, and give recommendations when each should be applied. We discuss the effect of unobserved species and the potential use of a threshold of minimum abundance for species inclusion. Finally, we consider these advances in the context of some of the main issues surrounding estimation of interaction sampling completeness in network ecology.

**Conclusions:** We recommend that future studies of bipartite networks utilise our approach and methods to estimate the sampling completeness of interactions, to assist with the quantitative and comparative analysis and interpretation of network properties.

## Background

The quantitative analysis of ecological networks can be directly affected by the proportion of all interactions that have been observed. This can be expected to increase with greater sampling effort in the field and more efficient means of detecting interactions in the lab [1–5], but will also vary by sampling method, by site or over time. This potentially compromises the comparability of ecological network analysis both within and between studies. Recent studies have attempted to address this by quantifying the proportion of interactions present in a system that have been sampled, using asymptotic species richness estimators [6–8]. The sampling completeness of species’ interactions may be used to confirm the validity of network analyses by checking that sampling is sufficiently ‘complete’, often defined, as a rule of thumb, as the detection of 90% of the interactions present. This has been proposed to balance adequate representation of the system against the diminishing return on effort when attempting to attain greater sampling completeness [6,9]. Sampling completeness can also be used to check for differences in sampling between different treatments in studies of replicated networks, which could potentially bias network metrics. Here, we review current methods to estimate sampling completeness and then propose an adaptation which should lead to improved estimates.

Chacoff *et al.* [6] were the first to propose estimating the sampling completeness of interactions in ecological networks. They recorded the occurrence of plant-pollinator interactions by observing flower visitation in 2048 separate samples of the study system. From this occurrence dataset they presented three estimates of sampling completeness in a bipartite mutualistic network: 1) sampling completeness of pollinator species alone (i.e. excluding interaction information); 2) sampling completeness of interactions for the whole network, based on the accumulation of plant-pollinator interactions across multiple samples of flower-visitor observations, and 3) sampling completeness of interactions for each plant species separately. In this latter case sampling completeness was estimated only for plant species with a minimum of 10 samples of flower-visitor observations and 10 observed visits by pollinating insects. In each method, the total abundance of species or interactions was estimated, using the Chao2 incidence-based estimator [10]. Sampling completeness was then calculated as the percentage of total species or interactions that were observed.

Although Chacoff *et al.*’s [6] whole-network estimate for sampling completeness of interactions has considerable merit, it is not universally applicable to all studies of bipartite networks, because the ‘incidence-based estimation’ depends on having multiple samples recording the detection (or not) of interactions. Importantly, sufficient independent samples are required to apply this approach (Chacoff *et al.* had 2048 discrete 5-minute observations of flower visitation, and 38 plant species observed more than 10 times [6]) and so it is possible to have too few samples if the taking of each discrete sample is labour-intensive [e.g. 7], even where overall sampling effort is high. One alternative is to use the Chao1 abundance-based estimator to directly estimate the total number of interactions based on the relative frequencies of unique interactions [7]; however, this may be inaccurate if the sample size is small [11], potentially resulting in sampling completeness being overestimated for the smallest (and therefore, potentially, the least complete) samples.

Traveset *et al.* [8], in a study of bird-flower visitation networks, had even greater sampling effort (∼500 hours of observations) than Chacoff *et al.* [6], but their observations were not so clearly partitioned into discrete samples. Therefore, they instead estimated the sampling completeness of interactions for the whole network by calculating sampling completeness of interactions for each species of flower-visiting birds as described above (retaining the minimum threshold of 10 individuals sampled for a species to be included), and then averaged across species using the arithmetic mean. Like Chacoff *et al.* [6], Traveset *et al.* [8] used the Chao2 estimator, but their method has the distinct advantage that it can be applied to networks generated by even a single sampling session (or multiple, aggregated sessions), treating each individual of a species as a discrete sample of that species’ interactions. Where the format of the data does not allow this approach, the Chao1 abundance-based estimator [12] could again be used to estimate the total number of interactions for each species based on the relative frequency of each interaction. For both estimators (but especially Chao1), this may be less accurate for locally-rare species (if the sample size is small); so this problem justifies the use of the minimum abundance threshold [8]. This approach is conditional upon the observed species in the focal level (i.e. it cannot consider the interactions of bird species that were not observed). The true sampling completeness (including the interactions of unobserved species) will therefore be lower than the estimated value, but by how much depends on the number of unobserved species and the number of their interactions, which will vary according to their (unobserved) identity. However, a benefit of this approach over that of Chacoff *et al.* [6] is that the species-level sampling completeness of interactions can be estimated directly from a bipartite network matrix in which columns contain species-level data for one level of the network, and rows contain individual-level data for the other level. Ultimately, the rows can be aggregated by species to construct the typical species-species interaction matrix in the format required for network analysis with standard software such as the R package bipartite [13]. By contrast, whole-network sampling completeness of interactions following Chacoff *et al.* [6] can only be calculated directly from an interaction matrix if an abundance-based estimator, such as Chao1, is used in place of Chao2.

However, by taking a simple *arithmetic mean*, the approach used by Traveset *et al.* [8] places equal weight on each observed species (not on each unobserved interaction), thereby placing proportionally more weight on the interactions of species that have few interactions (a small realised niche, whether because they are rare or because they are specialists). Here, we propose a modification of this approach that permits a more accurate estimation of sampling completeness of interactions within a network by taking a *weighted mean*, with each species weighted by its estimated interaction richness. Therefore we place equal weight on each interaction, whether or not it has been observed, rather than on each species. Our general approach is universally applicable to all studies of quantitative bipartite networks, through the use of two methods, which are selected depending on the nature of the sampling and resultant dataset (however, sampling completeness of interactions cannot be estimated for single binary bipartite networks using asymptotic species richness estimators).

In this paper we (i) introduce and describe our methods (*SC*_*W*_1 and *SC*_*W*_2), and discuss the scenarios in which each should be applied; (ii) demonstrate that our approach gives a mathematically accurate estimate of interaction sampling completeness, if all species of the focal level are observed; (iii) examine how sampling completeness varies if some species of the focal level are unobserved; and (iv) discuss some of the issues surrounding the estimation of interaction sampling completeness.

## Methods

### Description of our methods for estimating sampling completeness of interactions

Our approach is a weighted average version of that first used by Traveset *et al.* [8], but can be generalised to any quantitative bipartite network (Fig. 1). Interaction richness may be estimated using either of two methods, which respectively use abundance-based or incidence-based species richness estimators, depending on the nature of the sampling method used to detect interactions. Repeated sampling of interactions, either by taking multiple community-level samples or by sampling at the level of individuals, is not required to estimate sampling completeness of interactions using *SC*_*W*_1 (which applies an abundance-based estimator such as Chao1 [12]), but can be used to estimate sampling completeness of interactions more reliably using *SC*_*W*_2 (which applies an incidence-based estimator such as Chao2 [10]). In addition, our approach does not necessitate the use of a threshold for minimum number of individuals (as Traveset *et al*. [8] used), but can include all observed species. As specialist species tend to be rare and *vice versa* [14], this may reduce the risk of biasing the estimated interaction sampling completeness by primarily excluding specialists.

**Figure 1.**
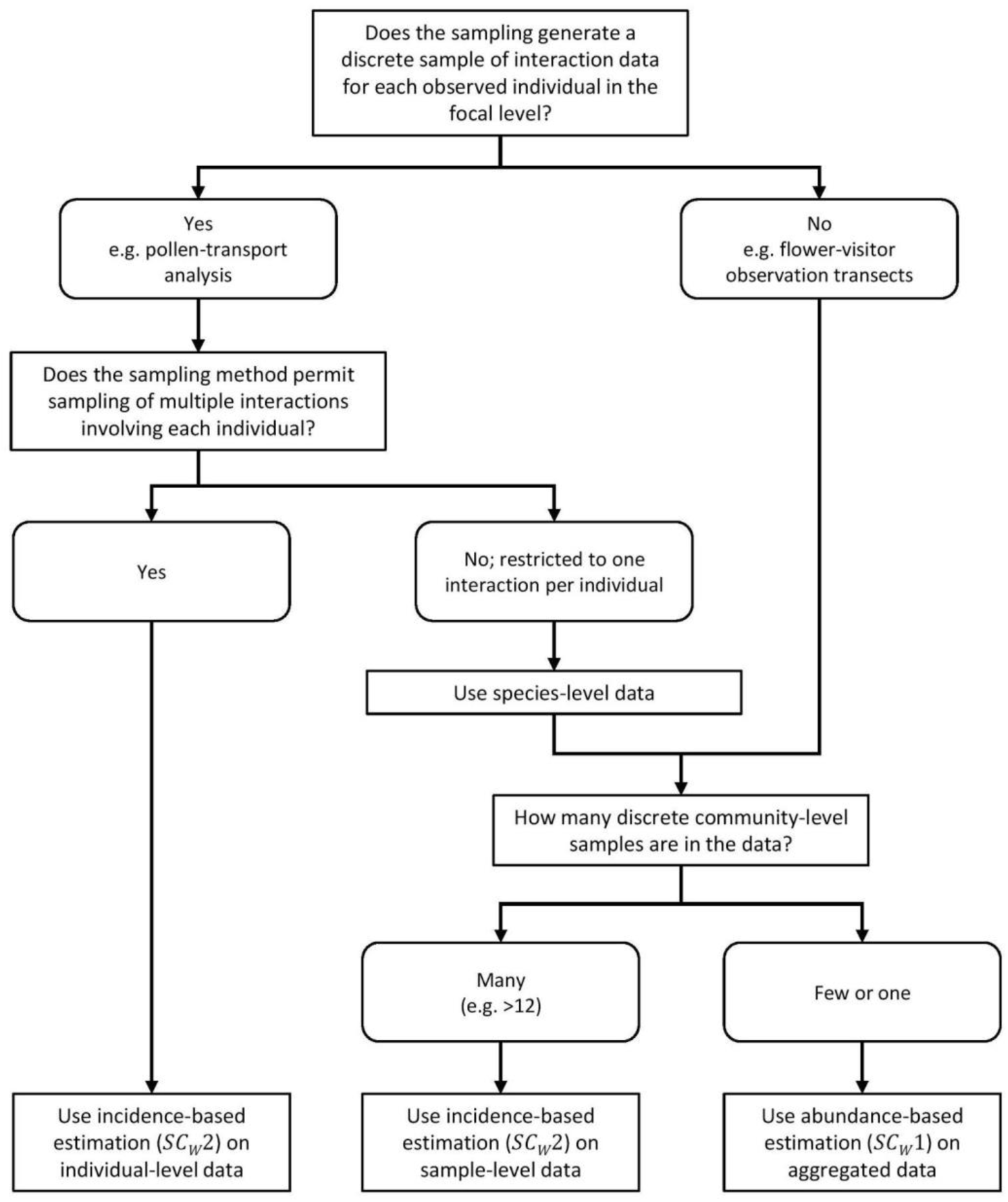
Flow diagram to determine which sampling completeness method to apply, depending on the nature of the dataset and of the sampling method used to detect interactions.

Throughout, we refer for simplicity to the approach used by Traveset *et al.* [8] as the unweighted sampling completeness (*SC*_*U*_), our proposed approach as the weighted sampling completeness (*SC*_*W*_), and the value which both methods attempt to estimate as the true sampling completeness (*SC*_*T*_). Additionally, we will refer generally to the calculation of *SC*_*W*_ which can be achieved by either of our methods, *SC*_*W*_1 (using abundance-based estimation) and *SC*_*W*_2 (using incidence-based estimation).

Bipartite ecological networks describe the interactions between two discrete levels of species; we refer to these as the “focal level” (the set of species on which observations were focussed; e.g. pollinators in pollen-transport analysis) and the “interacting level” (the set of species detected as a consequence of their interactions with the focal level; e.g. plants in pollen-transport analysis).

*SC*_*W*_1 uses interaction frequency of observed interactions (the number of individuals of a given species in the focal level observed to interact with a species in the interacting level) to estimate the number of unobserved interactions for each species in the focal level, using an abundance-based estimator such as Chao1 [12]. Interaction frequency has been shown to be a strong positive indicator of the strength of interspecific interactions [15], and can be readily generated for different interaction types, using various sampling methods. Therefore, *SC*_*W*_1 is applicable to any quantitative bipartite network, whether it is constructed from a single sample in which interactions are quantified or multiple samples that are aggregated to form a single network.

The *SC*_*W*_2 method may also be applied in studies where multiple discrete community-level samples are taken (e.g. multiple field surveys of a plant-pollinator community). In such cases *SC*_*W*_2 can be used to estimate the total interaction richness of each species in the focal level based on incidence of interactions involving that species in each sample. However, if the number of discrete community-level samples is small but sampling effort for each is high (leading to overall sample size being substantial), it may be more appropriate to use *SCW*1 on aggregated data from all samples than to use *SCW*2. Based on the performance analyses of Colwell and Coddington [11], we suggest that caution should be exercised if using *SCW*2 on fewer than 12 discrete samples.

Thus far we have considered use of the Chao1 and Chao2 estimators, but our approach could be applied using any species richness estimator. We have written generalized R functions to allow the estimation of sampling completeness for any suitable network using *SC*_*W*_1 and *SC*_*W*_2, and included these (along with a demonstration of their use) in Appendix S1. The R package vegan [18] permits species richness estimation using a range of estimators, and we have implemented all of these for use in our functions (Appendix S1). Specifically, with *SC*_*W*_1, it is possible to use either bias-corrected Chao1 [12,19] or ACE [19], whilst with *SC*_*W*_2, it is possible to use any of the bias-corrected Chao2 [10,19], first-order and second-order jack-knife [20], or bootstrap [21] estimators.

As it is based on species richness estimators, our approach assumes that the community is closed [10]. Our approach also assumes that the estimate of interaction richness computed using a species richness estimator approximates to the true interaction richness of each species. However, some generalist species may behave as specialists at the individual level [22]. In small samples, this has the potential to increase the ratio of singletons (interactions that appear in only one sample) to doubletons (interactions appearing in two samples), biasing the performance of species richness estimators towards a higher estimate of interaction richness, and lower sampling completeness. Therefore, the degree to which this assumption is true will depend on the level of similarity between individual-level and species-level specialisation. Nevertheless, we note that the same assumption is inherent in all previously-used approaches to estimating interaction sampling completeness, because they all utilise the Chao family of estimators.

### Mathematical justification of the weighted approach

#### Estimating sampling completeness, if all species in the focal level are observed

When estimating interaction sampling completeness by calculating the mean interaction sampling completeness of individual pollinator species, Traveset *et al*. [8] calculated the unweighted, arithmetic mean. However, the mathematical accuracy of this approach can be improved by weighting the mean by the estimated total number of interactions (interaction richness) of each species in the focal level (i.e. placing equal weight on each interaction, whether observed or not, rather than placing equal weight on each observed focal species). At the same level of sampling completeness, the absolute difference between estimated and observed interaction richness is greater for species which have many interactions (henceforth, “generalists”) than for those which have few (“specialists”). Therefore, an arithmetic mean of per-species sampling completeness may place undue weight on specialists, for which a relatively small number of unobserved interactions (making only a small contribution to network-level sampling completeness) can still lead to low species-level sampling completeness. Our approach allows a proportionally greater degree of weight to be apportioned to generalists than specialists when calculating the mean sampling completeness of all species.

We will demonstrate mathematically that, if all species in the focal level are observed, our approach equals the true value of sampling completeness.

Let:

*C* = percentage sampling completeness per species

*So* = observed interaction richness per species

*S*_*E*_ = estimated interaction richness per species

*S_T_* = true interaction richness per species

*n* = number of species *s*_*o*_

*m* = the subset of *n* species for which a minimum threshold number of individuals were sampled, where *m ≤ n*

Assuming that species richness estimators accurately estimate the true interaction richness (as stated above), then for a given species:

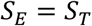

Percentage sampling completeness for each species is the percentage of the estimated interaction richness that has been observed:

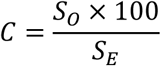

This can be arranged to:

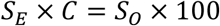

Likewise, the true sampling completeness of interactions is the percentage of the true interaction richness that has been observed, across all species:

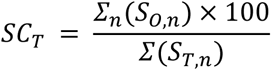

Our proposed approach estimates the sampling completeness of interactions by taking the mean sampling completeness per species, weighted by the estimated interaction richness:

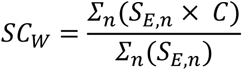

Drawing this together, it can be shown that our approach is mathematically equal to the true interaction sampling completeness when *E* is estimated accurately:

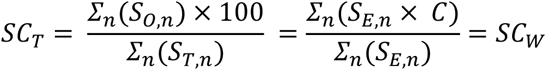

For comparison, the previously used partial, unweighted sampling completeness is not equal to the true sampling completeness, even for the subset of *m* observed species included in the estimate.

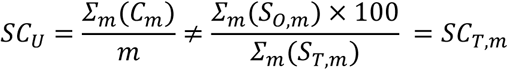

Therefore, if all species in the focal level are observed, our proposed approach will yield the true interaction sampling completeness.

#### Estimating sampling completeness, if some species in the focal level are unobserved

Both the approach of Traveset *et al*. [8] and our adjusted approach are conditional upon the observed set of species in the focal level. Although these approaches therefore allow the relative specialisation of each species to be taken into account, they also introduce the possibility of inaccuracy if some species in the focal level are unobserved; a scenario that is likely in the majority of studies of bipartite ecological networks.

In addition to the above, let:

*U* = cumulative interaction richness of all unobserved species in the focal level If all focal species are observed, this is 0, but otherwise it is positive:

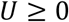

Because there are now unobserved species, with *U* interactions, of which zero are observed:

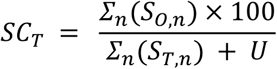

Because a fraction with the same numerator and a larger denominator must be smaller, we can infer that:

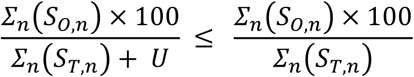

and, from above,

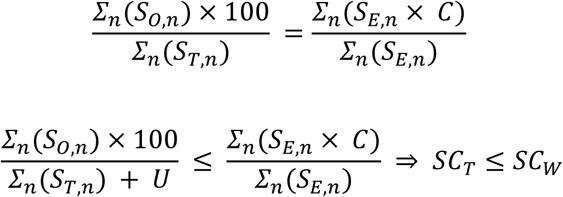

Therefore, if some species in the focal level are unobserved, our approach will always overestimate the sampling completeness of interactions. This allows us to state that the true sampling completeness of interactions for the whole network (including unobserved species) is “up to” the value estimated by our approach. The smaller the value of *U*, the closer the estimate of our approach will be to the true value of sampling completeness of interactions.

Our approach is therefore most accurate if unobserved species have a low number of interactions and make little contribution to the overall interaction richness of the network (so that their true weight is close to the weight of zero that they are effectively assigned). Crucially this assumption is ecologically reasonable, because unobserved species are likely to be rare, and rare species tend to be functionally specialist (even if their fundamental niche is generalist) [14]. It is therefore likely that most unobserved species will either be specialists or appear to be specialists.

## Results

To test and demonstrate the use of our approach through the methods *SC*_*W*_1 and *SC*_*W*_2, we used each method to estimate the sampling completeness of interactions for suitable interaction datasets. To demonstrate that *SC*_*W*_1 is universally applicable to all quantitative, bipartite networks, we downloaded all 16 empirical datasets included as examples in the R package bipartite [13] (Table S1). Each dataset represents a single quantitative plant-pollinator network. We estimated the sampling completeness of each network using *SC*_*W*_1 with both the Chao1 [12] and ACE [19] estimators (Fig. 2). We found that sampling completeness estimated using Chao1 ranged widely, from 35.6% (for the ‘kato1990’ [23] network) to 100% (for the ‘bezerra2009’ [24] and ‘olesen2002flores’ [25] networks). Besides these, only one other network met the 90% rule of thumb for sufficiently complete sampling (‘olesen2002aigrettes’ [25], 98.7% complete using *SC_W_*1 with Chao1). There was strongly significant positive correlation between the estimates of sampling completeness using Chao1 and using ACE (Pearson’s r, t = 16.15, d.f. = 14, p < 0.001).

**Figure 2.**
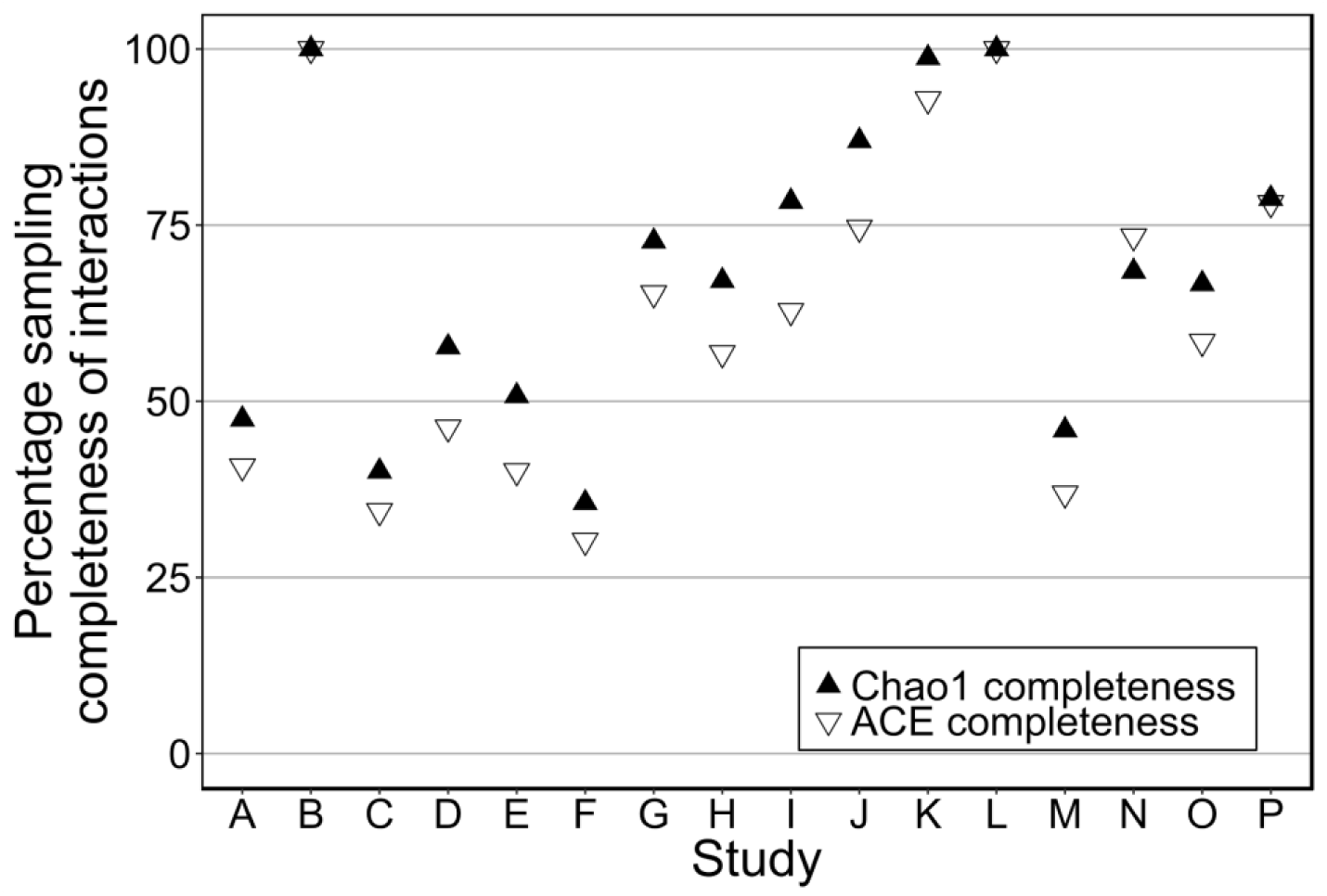
Estimated sampling completeness of interactions for 16 empirical plant-pollinator networks included in the R package bipartite [13]. Sampling completeness was estimated using *SC*_*W*_1 (abundance-based estimation), using both the Chao1 and ACE estimators. Citations to datasets shown are given in Table S1.

However, although all 16 networks included in bipartite [13] are quantitative, none include either individual-level data on the focal level, or data from discrete sampling sessions, so *SC*_*W*_2 cannot be used. Therefore, to demonstrate the use of *SC*_*W*_2, we used data from Macgregor *et al.* [26,27]. This dataset contains nocturnal plant-pollinator interactions observed by sampling pollen transport from the proboscides of individual moths (Lepidoptera), and the individual-level data on the focal level (moths) is retained, making it suitable for estimation by *SC*_*W*_2. We estimated the sampling completeness of the network using *SC*_*W*_2 with the Chao2 [10,19], first- and second-order jackknife [20], and bootstrap [21] estimators and, for comparison, we also estimated sampling completeness of the same network using *SC*_*W*_1 and the Chao1 and ACE estimators (Fig. 3). Sampling completeness was generally estimated to be around 60% when using all of the Chao2 (57.2%), first-order jackknife (65.7%) and second-order jackknife (54.6%) incidence-based estimators and both the ACE (59.9%) and Chao1 (66.0%) abundance-based estimators, but was estimated to be substantially higher (81.5%) when using the bootstrap estimator.

Using the same network, we tested the impact of a threshold minimum number of individuals for a species’ inclusion (as applied by Chacoff *et al.* [6] and Traveset *et al.* [8]) on the estimation of sampling completeness using *SC*_*W*_2. We estimated sampling completeness of interactions for every threshold level between 1 (all observed species retained) and 10 (all species with fewer than 10 observed individuals excluded), using *SC*_*W*_2 with the Chao2 estimator (we chose Chao2 for this test because it is the most robust estimator to small sample sizes [11]). We found that the number of species included in the sampling completeness estimate decreased from the total of 202 observed species to only 35 when the 10-individual threshold was applied, and that estimated sampling completeness changed unpredictably depending on the level at which the threshold was set (Fig. 4). In general, higher thresholds led to lower estimates of sampling completeness, but an increase in sampling completeness between thresholds of 7 and 8 individuals demonstrated that the level at which the threshold is set is arbitrary. Nevertheless, sampling completeness was estimated to be highest when all species were retained (57.2%) and lowest when all species with fewer than 10 individuals were excluded (49.8%).

## Discussion

### Issues surrounding the estimation of interaction sampling completeness

#### Threshold for minimum number of individuals

When estimating species-level sampling completeness of interactions, both Chacoff *et al.* [6] and Traveset *et al.* [8] included only species for which at least 10 individuals had been sampled. The accuracy and precision of species richness estimators decreases for small samples [28], and so the use of this threshold is intended to ensure that the sampling completeness of interactions is calculated from only the most accurate estimates of interaction richness, even at the expense of some biological information. Although we have implemented the option for such a threshold in the R code that accompanies this paper (Appendix S1), we nevertheless prefer not to apply such a threshold with our approach (specifically with *SC*_*W*_2), for several reasons.

Firstly, such a threshold would not be universally applicable for *SC*_*W*_1, and might therefore lead to discrepancies in the estimation of sampling completeness between *SC*_*W*_1 and *SC*_*W*_2. Our further arguments therefore refer specifically to the application of a threshold when using *SC*_*W*_2.

Secondly, the number of individuals at which the threshold is set is arbitrary, and the final estimate of sampling completeness will vary unpredictably depending on the chosen threshold (Fig. 4). Additionally, exclusion of rare species (i.e. those with few individuals) by applying a threshold could lead to overestimation of sampling completeness, because these species would effectively be treated as if unobserved. Because specialist species are more likely to be rare [14], they are more likely to be excluded by the application of a threshold; this could potentially introduce further bias to the estimated sampling completeness.

By the same logic, because rare species (within the study system) are more likely to be functionally specialist, they are likely to be accorded low weight and therefore any inaccuracy in the estimation of interaction richness for these species will have little impact on the final estimated value of sampling completeness, reducing the need for their exclusion. This may be further assisted by the use of the Chao2 estimator, which is one of the least biased species richness estimators for small numbers of samples [11], and so may minimise the potential for such inaccuracy. Additionally, because the Chao2 estimator technically provides the lower bound for species (or interaction) richness [10], it is more likely to underestimate richness than overestimate it [e.g. 29]. As a result, any inaccuracy in estimation for species with few individuals is likely to lead to lower weight being assigned to those species when calculating the final estimate of sampling completeness.

Therefore, the use (or not) of such a threshold represents a trade-off between the error introduced by including low-abundance species (for which interaction richness may not be accurately estimated) and the error introduced by treating such species as if unobserved. However, because species sampled at low abundance are likely to be relatively specialist (and therefore assigned low weight, if our approach is used), we believe that their inclusion in estimation of sampling completeness is relatively safe. Given this, we also believe that it is more appropriate to include all species, due to the potential to introduce bias to the estimated sampling completeness of interaction by treating rare species as if they are unobserved.

#### Deciding on the focal level - upon what is the sampling completeness conditional?

Our estimate of sampling completeness, like that of Traveset *et al.* [8], is conditional on one of the levels of the bipartite network (referred to here as the focal level). In other words, as justified in the previous sections, the estimate assumes that the focal level has been completely sampled. Our estimate is of the maximum sampling completeness, and the true value (i.e. including unobserved focal species) is less than or equal to this estimate. If the number of interactions with unobserved species in the focal level is small, then the estimate of sampling completeness is not much less than the true value.

The definition of the focal level is that it is directly constrained by the sampling, whereas the interacting level is directly constrained by the focal level. Often the focal level is obvious: for example, birds producing seed-filled droppings or insect pollinators transporting pollen grains. In these cases it is clear that the individual animal is directly sampled, and the identity of seeds in droppings or pollen on insects depends upon the preceding behaviour of the individual. Focal observations of flowers are similar, with the plant being the focal level. Other situations are less obvious, most notably plant-pollinator transects where individual insects are sampled whilst visiting flowers. In this example we suggest that the plants should be viewed as the focal level because, in theory, the sampling is constrained by the plants that are present, whereas the insect pollinators are mobile and their presence is dependent on the flowers present in the transect.

#### Choosing between SC_*W*_1 and SC_*W*_2

We have discussed the situations in which *SC*_*W*_1 and *SC*_*W*_2 can be applied in the descriptions of each method, but here we will synthesize the process of deciding between the two (Fig. 1). Although *SC*_*W*_1 can be applied to any quantitative bipartite network, we recommend using *SC*_*W*_2 where appropriate, due to the greater robustness of the Chao2 estimator to the effects of small sample sizes [11]. The first consideration should be whether it is possible to independently sample the interactions of each individual in the focal level, and if so, whether it is possible to sample multiple interactions from a single individual. If the answer to both questions is yes (e.g. sampling seeds from the droppings of birds, where each dropping can be linked to the individual bird from which it was sampled, and multiple seeds can be detected in each dropping), then *SC*_*W*_2 can be used. However, if it is only possible to sample a single interaction per individual (e.g. sampling host-parasitoid interactions by rearing, where it is only possible for a single parasitoid to emerge from each host), it may be more appropriate to aggregate the data at species-level, as the level of generalisation will differ at individual- and species-level: all individuals will appear to be extreme specialists even if the species is generalist.

If, however, the network data does not allow the assessment of individuals in the focal level - either because the sampling methods do not permit the collection of such data (e.g. flower-visitor transects where the species of plants in the focal level are collected, but not the identity of each individual plant) or because such data have been aggregated at species-level, then it may still be possible to apply *SC*_*W*_2 by examining incidence across multiple samples of the network, depending on the number of discrete samples that have been taken. Overall sample size may be large even if the number of discrete samples is small, depending on the effort invested in obtaining each discrete sample. Performance analyses by Colwell & Coddington [11] suggest that the Chao2 estimator accurately estimates the true number of entities when the number of samples is 12 or more. Therefore, for fewer than 12 samples we recommend using *SC*_*W*_1on pooled data in order to maximise the effective sample size.

#### Assessing the influence of unobserved species in the focal level

As we have previously discussed, the interactions of unobserved species belonging to the focal level will have an unknown influence upon the true value of sampling completeness. It is possible to assess the likely influence of unobserved focal species, in the cases where species of the focal level are sampled in proportion to their abundance (either as part of the sampling of interactions, or in addition to it). Asymptotic species richness estimators can be used to estimate the number of unobserved species in the focal level. If the number of unobserved species in the focal level is small, then these are likely to have little impact on sampling completeness, therefore that the estimate of *SC*_*W*_ will not be very different from *SC*_*T*_.This is especially the case if the unobserved species are functionally relatively specialist, which is to be expected if their abundance is low.

#### Considering uncertainty of the estimate

Throughout we have considered the point estimate of sampling completeness and have not included its uncertainty. The question of how accurately community-level sampling completeness can be estimated is nonetheless important. Given that species richness estimators often have high uncertainty, the uncertainty of sampling completeness is likely to be considerable. This variation is further increased by the possibility of choosing any of several species richness estimators, which may differ in their estimated interaction richness for each species (Fig. 3). Here we briefly discuss several methods that would typically be used to estimate uncertainty around a point estimate and why they are not suitable for sampling completeness.

**Figure 3.**
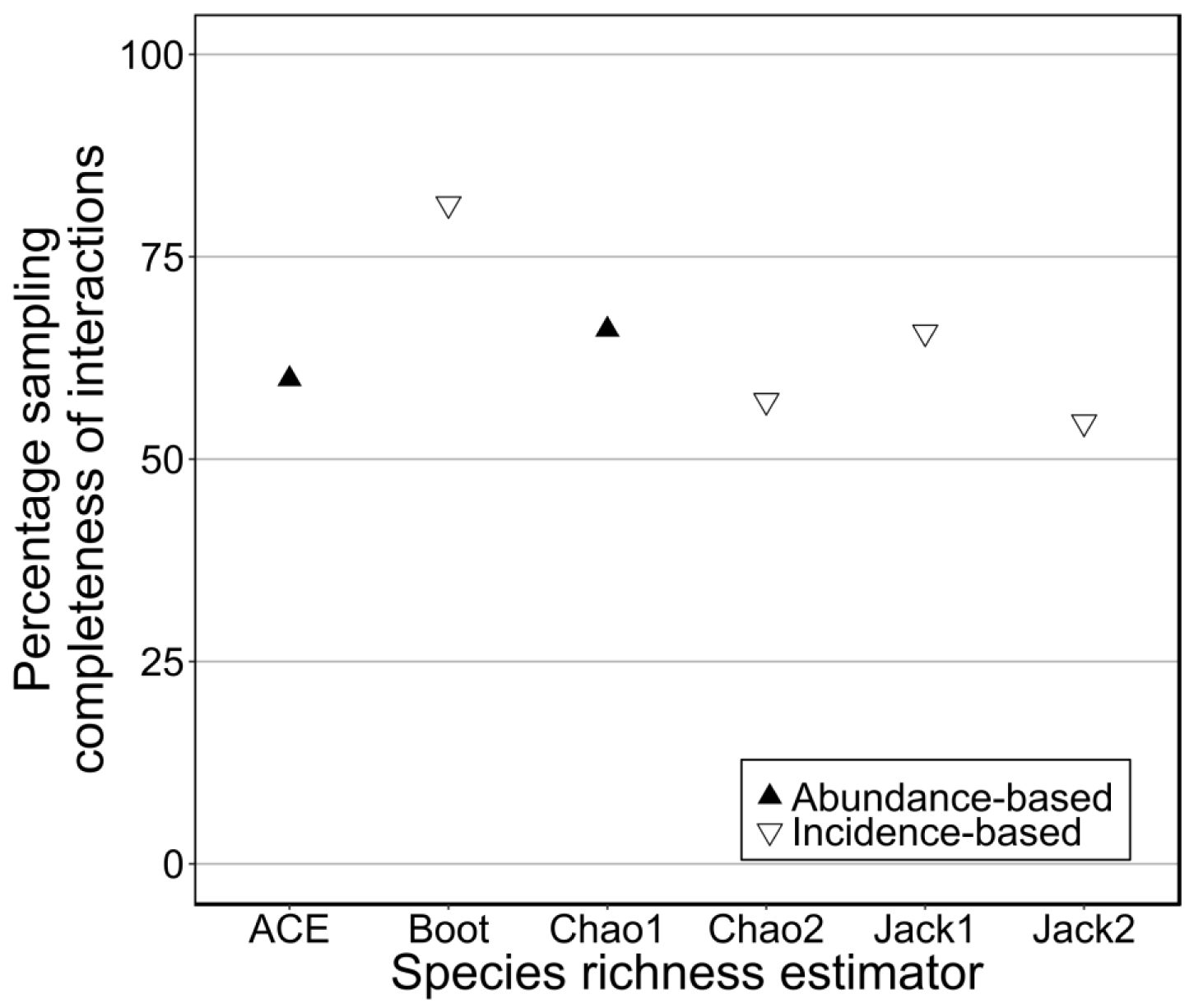
Estimated sampling completeness of interactions for an empirical plant-pollinator network [26,27] calculated using *SC*_*W*_1 (abundance_-_based) and *SC*_*W*_2 (incidence-based), with a range of estimators. Sampling completeness was calculated using the ACE, bootstrap (“Boot”), Chao1, Chao2, first-order jackknife (“Jack1”) and second-order jackknife (“Jack2”) species richness estimators; black triangles indicate abundance-based estimators and white triangles indicate incidence-based estimators.

**Figure 4.**
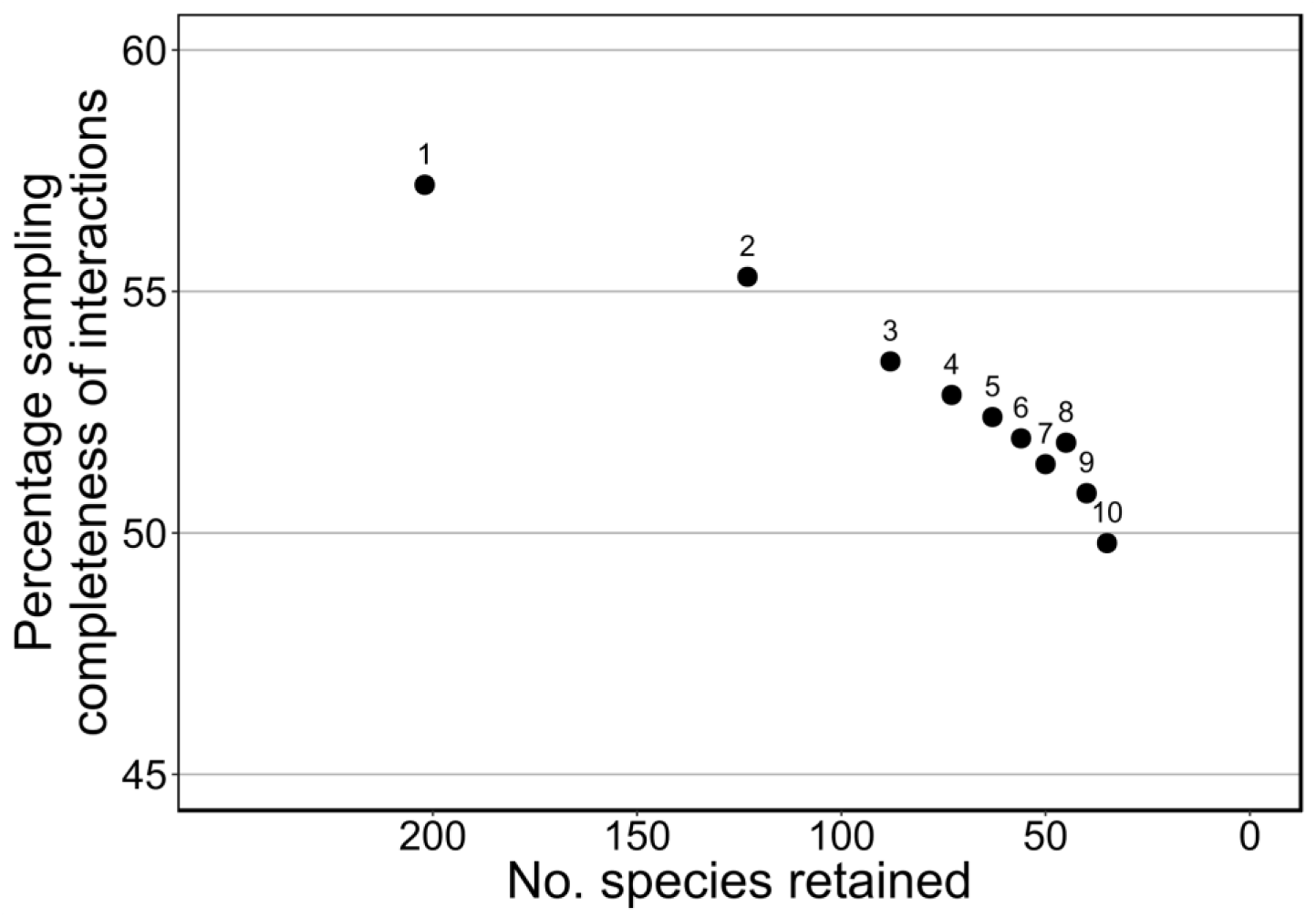
Estimated sampling completeness of interactions for an empirical plant-pollinator network varies unpredictably according to the level at which we set the threshold for minimum number of individuals for a species to be included. Sampling completeness was estimated at different threshold levels from 1 individual (all species retained) to 10 individuals (per Traveset *et al.* [8]). Points are labelled with their threshold level and show the number of species retained (out of a total of 202 observed species) and the estimated sampling completeness of interactions.

Species richness estimators, including both Chao1 and Chao2, provide a point estimate of the number of unobserved entities (when calculating sampling completeness, these entities are interactions), the variance of which is normally distributed around the log-transformed estimate. This is added to the number of observed entities to give the estimated true number of entities, so *S*_*E*_ = *S*_*0*_ + *S*_*U*_ where *log*(*S*_*U*_) *∼ N*(*μ,σ*). This in turn forms the denominator in the per-species sampling completeness (*SC*_*W*_= *S*_*O*_/*S*_*E*_). Mathematical operations can be undertaken on the variance of distributions, but as the variance on the log-scale forms part of the sum of the *denominator*, it is effectively intractable to carry through mathematically.

An alternative approach is to use randomisation and we considered two ways to do this. Firstly, we considered Monte Carlo resampling of the variance of the estimates. Sampling from *log*(*S*_*U*_) *∼ N*(*μ,σ*) creates a distribution, but the inverse logarithm results in a highly skewed distribution of *S_U,rand_* and hence exceedingly large values of *S_E,rand_ = S_O_ + S_U,rand_*. High values of *S_E,rand_* have a dual effect: (i) sampling completeness will be low because *S/_O_ S_E,rand_, S_O_* in is fixed, and (ii) high weight will be given to species with high values of *S_E,rand_* when calculating a weighted average across species. A system that gives disproportionately high weight to species with disproportionately low sampling completeness will produce an overall sampling completeness is lower than expected. So, although variance of randomised sampling completeness can be calculated, its mean will be biased low.

Secondly, we could resample the raw data and there are two ways of doing this: bootstrapping and creating null models. Bootstrapping involves resampling interactions with replacement and is a widely used method to obtain estimates of variance of metrics. So, interactions could be sampled (with replacement) from the observed set of interactions to create a new, random matrix of interactions to give *S*_*O,rand*_ for each species. An equivalent way of achieving the same would be to randomly choose interactions, up to a certain sample size, according to their relative proportions in the raw data. As before, we can calculate *S*_*E*_ using the appropriate estimator, and hence *SC*_*W*_, and could repeat this many times to calculate variance. However, *So*_*rand*_ is constrained: *So* _*rand*_ for a species could be less than observed in the raw data, but it could never be more (just as when randomly choosing, with replacement, beads from a bag of black and white beads, a sample could comprise one or two colours of beads, but never three). Bootstrapping should have no bias on S_o,rand_ because it is an estimate based on a sample (whether the raw data or the random sample from the raw data), although it might affect its precision. However, if *S*_*o,rand*_ is biased low, then *SC*_*W*_*= So* _*rand*_ */S*_*E*_ will also be biased low.

The second way of resampling the raw data is to create a null network based on redistributing interactions within the network according to particular constraints (e.g. constraining the row and column sums, and/or the network connectance, using functions such as swap.web or vaznull from the R package bipartite [13]). The resulting network will be a result of the null models and even for highly conservative models, they assume that species associate randomly. They therefore tend to increase the degree to which species in the network appear to be generalists [30,31], and reduce the occurrence of singletons in the network relatively more than they reduce the occurrence of doubletons. As singletons form the numerator in the majority of species-richness estimators [32], this leads to systematically smaller estimates of true interaction richness, and because the number of observed interactions is fixed sampling completeness is biased high.

Overall, estimates of the precision of sampling completeness would assist with its proper interpretation, but currently these are not currently obtainable in an unbiased way. This would be a valuable direction for future investigation.

### Realised vs fundamental niche

Our approach estimates the sampling completeness of the realised niches of each species in the network, rather than their fundamental niches. This distinction is most simply explained in the context of rare generalist species, which might have the ecological potential to interact with a wide range of species in the network (and throughout their global range may indeed do so), but in practice only interact with a subset of those species in the system under study, because each individual interacts independently with a subset of its fundamental niche, and there are few individuals. Estimating the sampling completeness of the realised niche is appropriate, because a potential interaction that is not realised, by definition, cannot be sampled. Failure to sample such an interaction therefore does not indicate incomplete sampling. Nevertheless, it could be of interest in some studies to estimate the proportion of potential interactions that are realised. In such cases, an approach based on forbidden links (interactions that never occur, and which are therefore outside a species’ fundamental niche) may be more appropriate [see 33,34].

### Developing sampling techniques

Although *SC*_*W*_1 could be applied to any quantitative bipartite network, *SC*_*W*_2 requires either individual-level data on the focal level, or networks constructed from many repeated sampling events for each focal-level species. However, most previous empirical studies of ecological networks have focussed on small numbers of networks that aggregate data from multiple sampling sessions [e.g. 35]. Many common methods for sampling interactions do not permit generation of suitable individual-level data. For example, flower-visitor observation transects [e.g. 36] generally do not collect individual-level data about the focal level (plants), whilst although host-parasitoid rearing collects individual-level data, it cannot detect more than one interaction per individual, even if multiple interactions exist [37]. Recent developments in DNA-based approaches to detecting and identifying interspecific interactions (such as DNA metabarcoding) offer considerable potential to increase the scale and resolution of data collection in ecological network analysis [38]. Where this is based on obtaining multiple interactions per sample, e.g. pollen on insects or faecal remains, data collected by such approaches are likely to be well-suited to estimation of interaction sampling completeness using *SC*_*W*_2, because DNA extraction methods tend to focus on individuals of the focal level. DNA metabarcoding may also facilitate the detection of multiple interactions per individual for interaction types where current sampling methods do not permit this, such as host-parasitoid interactions [37,39].

## Conclusions

Estimating sampling completeness is important because of its influence on descriptive network metrics. Our proposed approach for estimating the sampling completeness of interactions in quantitative bipartite networks is to calculate the weighted mean of the sampling completeness calculated for all observed species in the focal level. This builds upon the approach used by Traveset *et al*. [8], increasing its mathematical accuracy by reducing the influence of species with few interactions, and carries several advantages over the previously-used approaches. We show the difference between incidence-based and abundance-based methods and discuss when each method is appropriate. We show that further research is necessary to obtain measures of precision for estimates of sampling completeness, and that this would be valuable to the interpretation of sampling completeness estimates.

We recommend that future studies of bipartite networks estimate the sampling completeness of interactions by taking a mean of the estimated interaction sampling completeness of all focal species, weighted by the estimated interaction richness per species, and use this estimate to help interpret differences when undertaking comparative analyses of networks.

## Declarations

### Ethics approval and consent to participate

Not applicable

### Consent for publication

Not applicable

### Availability of data and material

Datasets analysed in the production of Fig. 2 are included in the R package bipartite [13]; full citations are given in Table S1. The dataset analysed in Figs. 3 & 4 is from Macgregor *et al.* [26,27] and can be accessed at https://doi.org/10.5285/31cc5cec-d33b-4dd6-a932-061ff947e708

### Authors’ contributions

C.J.M. initially conceived the idea of weighted sampling completeness (*SC*_*W*_). All authors contributed to developing the methods *SC*_*W*_1 and *SC*_*W*_2 and preparing the manuscript.

## Acknowledgements

We are grateful to J. Tylianakis and R. Sanderson for their useful feedback on an early version of the manuscript.

## Supplementary files

### Appendix S1

R scripts and example data to demonstrate the application of *SC*_*W*_1 and *SC*_*W*_2. Contains:

#### Appendix S1.1

An RMarkdown file demonstrating the application of *SC*_*W*_1 and *SC*_*W*_2 through a series of worked examples (including the creation and analysis of Figs. 2-4).

#### Appendix S1.2

An R script containing fully-generalised functions for the calculation of *SC*_*W*_1 and *SC*_*W*_2 on any suitable dataset. This script is sourced in Appendix S1.1.

#### Appendix S1.3

An empirical dataset used for the demonstration of *SC*_*W*_2 in Appendix S1.1. This dataset has been pre-processed into the correct format for applying *SC*_*W*_2; the original, unprocessed dataset is available online [26].

***Table S1***

List of citations to example datasets included in the R package bipartite and analysed in the production of Fig. 2.

